# Deletion of the polarity protein Llgl1 impairs cardiomyocyte intercalated disc integrity and permits cell cycle activity into adulthood

**DOI:** 10.64898/2025.12.19.695635

**Authors:** Jerrell J. Lovett, Michael A. Flinn, Amirala Bakhshian Nik, Anooj Arkatkar, Caelan W. Moreno, Caroline J. Greenheck, Sydney L. Buday, Kaelin A. Akins, Brianna R. Waack, Julia N. Pindaro, Alexandra L. Purdy, Siqi Li, Bradley S. Miller, Lu Han, Chun Liu, Michaela Patterson, Brian A. Link, Caitlin C. O’Meara

## Abstract

**Background:** Cell polarity orchestrates spatial organization within tissues influencing cellular processes such as adhesion, proliferation, and differentiation. While the role of polarity proteins is well-established in epithelial systems, their function in mammalian cardiomyocytes remains poorly understood. Global deletion of the polarity protein, Llgl1, has been shown to impair cardiac trabeculation and disrupt cardiomyocyte junctions in zebrafish. Here, we investigate the role of Llgl1 in mammalian cardiomyocyte function.

**Methods:** We developed a mouse model whereby Llgl1 is deleted from cardiomyocytes (Llgl1^CKO^) and assessed cardiomyocyte intercalated disc integrity, cell cycle activity, cardiac function, and electrophysiology. We subjected mice to myocardial infarction to test if permanent Llgl1 deletion, or inducible Llgl1 deletion in adult cardiomyocytes (Llgl1^TamCKO^), influences the cardiac injury response. We performed RNAsequencing and in vitro knockdown and viral overexpression experiments to delineate the mechanisms by which Llgl1 influences cardiomyocyte phenotypes.

**Results:** Llgl1^CKO^ mice display disrupted intercalated disc integrity indicated by reduced protein density at cardiomyocyte junctions, associated with electrical conduction defects that manifest as ventricular arrhythmia. Furthermore, Llgl1^CKO^ mice display enhanced cardiomyocyte cell cycle activity extending into adulthood. Llgl1^CKO^ mice and mice whereby Llgl1 is deleted by tamoxifen induction in adulthood both show improved cardiac function and reduced scar size following myocardial infarction. Mechanistically, Llgl1 alters mechanotransduction and cell cycle pathways in the heart, and regulates proper localization of N-cadherin to the cell membrane, a process that is optimized by phosphorylation of five serine residues at the C-terminus of Llgl1.

**Conclusions:** We elucidate a key role for Llgl1 in establishing proper intercalated disc integrity in mammalian cardiomyocytes. Disruption of intercalated discs and mechanotransduction via Llgl1 knockout is associated with cardiac arrhythmia, elevated cell cycle activity and resistance to cardiac injury. These studies shed light on mechanisms underlying intercalated disc establishment and connect disrupted cardiomyocyte junctional integrity with enhanced cell cycle activity.

## INTRODUCTION

Cell polarity is a fundamental and evolutionarily conserved process crucial for establishing spatial organization within cells and tissues. It governs essential cell behaviors, including fate determination, morphology, proliferation, and adhesion, by directing the asymmetric distribution of proteins and other macromolecules across cellular domains.^1,2^ Within epithelial cells, disruption of polarity factors leads to defects in cell junction formation and in some cases permits uncontrolled cell proliferation, as observed in *Drosophila* and vertebrate models.^3–5^ Despite a wealth of data on polarity factors in epithelial systems, the role of these proteins in cardiomyocyte (CM) biology and cardiac function postnatally remains poorly understood.

The apical-basal cell polarity factor, Lethal (2) giant larvae protein homolog 1 (Llgl1) has been well described for its roles in regulating polarity in epithelial cells. Llgl1 regulates the localization of atypical protein kinase C (aPKC) to the apical Par complex by competitive interaction with partitioning-defective-6 (Par6).^6^ Reciprocally, aPKC phosphorylates Llgl1 to exclude it from the cell apical domain^7^ and this mutual antagonism is a key mechanism by which epithelial cells establish classical apical-basal polarity.^8^ Components of the Scribble (*scrib*) polarity complex comprised of *scrib*, discs-large (*dlg*) and *lgl*, act together to regulate cell polarity and cell-cell adhesion. Mutation of any of these three proteins disrupts adherens junctions in epithelial cells which is accompanied by over-proliferation phenotypes.^3^ Llgl1 is known for its tumor-suppressive properties across multiple tissues as mutation of the *Drosophila* homologue, *lgl*, results in tumor formation of imaginal discs and neuroblasts,^9^ and in humans Llgl1 expression inversely correlates with aggressiveness in cancers such as squamous lung carcinoma, breast cancer, and pancreatic cancer.^10–12^

Llgl1 stabilizes cell junction complexes in epithelial cells as well as non-epithelial cell types such as mouse neural stem cells. In neural stem cells, Llgl1 expression mediates N-cadherin localization to the cell membrane, a process dependent on Llgl1 phosphorylation by aPKC.^6^ In many cell types including CMs, N-cadherin is the primary mediator of cell-cell adhesion, forming homodimers with neighboring cells to create a zipper-like intercellular structure.^13^ N-cadherin is essential for cardiac development, as its loss results in global disruption of the intercalated disc (ICD); specialized junctions that facilitate cell-cell communication and contraction synchronization,^14–17^ leading to sudden cardiac death.^18,19^ These observations highlight the importance of proper N-cadherin localization in initiating ICD establishment. However, the comprehensive mechanisms that govern ICD establishment and protein accumulation at the termini in CMs remain unclear.

While the role of Llgl1 in mammalian CMs has not been directly explored, global deletion of *llgl1* in zebrafish disrupts cardiac development. Myocardial trabeculation is facilitated by basal cell delamination and in zebrafish *llgl1* is required for basal laminin remodeling during trabeculation.^20^ In addition to trabeculation defects, *llgl1* global knockout zebrafish display dysmorphic CM ICDs characterized by protein mis-localization and expanded ICD domains.^20,21^ Studies in zebrafish collectively implicate a role for Llgl1 in heart development and CM biology, however, these prior studies used global knockout zebrafish models therefore the intrinsic role for Llgl1 in CMs, particularly in mammalian CMs, remains unknown.

Considering studies in zebrafish implicate a role for Llgl1 in CM biology, we hypothesized that Llgl1 may play a pivotal role in establishing or maintaining CM cell junctions and potentially influencing cardiac physiology and CM cell cycle activity in mammalian systems. In the present study, we demonstrate that Llgl1 deletion specifically in mouse CMs (Llgl1^CKO^) is required for establishing proper composition of ICDs, a phenotype observed into adulthood. Disrupted ICDs is associated with ventricular arrhythmia in adult Llgl1^CKO^ mice and, coincidentally, with increased CM cell cycle activity in both uninjured settings and following myocardial infarction (MI) in adults. Llgl1^CKO^ mice as well as tamoxifen mediated induction of Llgl1 deletion in adult CMs (Llgl1^TamCKO^) improves ejection fraction and reduces scar size following injury. Llgl1 deletion is associated with modulation of cell cycle genes, integrins, and mechotransduction pathways in sham operated and post MI hearts, suggesting that altering cell cycle or mechanotransduction via Llgl1 depletion protects CMs during the injury response.

Mechanistically, we show that like neuronal epithelial cells Llgl1 mediates proper membrane localization of N-cadherin in CMs. N-cadherin membrane localization in CMs is in part dependent on phosphorylation of Llgl1 at five serine residues at its C terminus (S982A, S986A, S988A, S989A, S992A); residues previously reported to be differentially phosphorylation during neonatal mouse heart regeneration.^22^ The serine residues mutated in our study are distinct from the previously reported aPKC target residues that regulate Llgl1 mediated N-cadherin localization in neuronal epithelial cells suggesting a distinct mechanism of Llgl1 regulation.^6^ Collectively, these findings elucidate Llgl1 as a key regulator in establishing membrane localization of ICD proteins and loss of Llgl1 impairs proper CM junctional complexes which is associated with arrhythmia, protection from adverse ventricular remodeling, and permissive CM cell cycle activity into adulthood.

## METHODS

### Animals

Mice were housed in the Biomedical Resource Center at the Medical College of Wisconsin, an AAALAC-approved facility. All animal protocols were approved by the local IACUC (AUA 00005516). Llgl1 floxed (Llgl1^fl/fl^) mice^6,23^ were a generous gift from Dr. Valeri Vasioukhin at the Fred Hutchinson Cancer Research Center, Seattle Washington. Llgl1^fl/fl^ mice were crossed with B6.FVB-Tg(Myh6-cre)1282Mds/J mice ([Myh6^Cre^] Jackson Laboratories 011038) to generate Llgl1^fl/fl^;Myh6^Cre^ (“Llgl1^CKO^”) mice and Llgl1^wt/wt^;Myh6^Cre^ littermate controls (“Myh6^Cre^”). Llgl1^fl/fl^ mice were also crossed with B6.FVB(129)-A1cf^Tg(Myh6-cre/Esr1*)1Jmk^/J ([Myh6^MCM^] Jackson Labs 005657) mice to generate Llgl1^fl/fl^;Myh6^MCM^ (“Llgl1^TamCKO^”) and Llgl1^wt/wt^;Myh6^MCM^ littermate controls. For all *in vivo* mouse experiments both males and females were used and data from both sexes were pooled. Pregnant Sprague Dawley rats were purchased from Charles Rivers Laboratories (strain #400) and postnatal day 1 (P1) Sprague Dawley rat pups were used for neonatal ventricular CM (NRVCM) isolation and culture.

### 5-Ehtynyl-2’-deoxyuridine (EdU) administration

For neonatal experiments, EdU was administered by subcutaneous injection at a dose of 1 mg/kg at P4 and P5. Hearts were then collected for analysis of EdU incorporation at P7. For adult post MI experiments, animals were injected with 10 mg/kg EdU by intraperitoneal injection daily from 4 to 10 days post injury (dpi). For osmotic minipump experiments, minipumps (Alzet, model 1007D) were loaded with 2.5 mgs of EdU which was diluted to 100 mg/mL in DMSO and then further diluted to 2.5 mg/100 µl in sterile saline. EdU loaded minipumps were implanted subcutaneously at 4 dpi. Pumps were removed at 10 dpi and hearts were collected the following day for CM nucleation and ploidy analysis.

### Genomic DNA PCR

Lysis buffer (2 M Tris pH 8.5, 0.5 M EDTA pH 8.0, 20% SDS, 5 M NaCl, sterile autoclaved dH_2_O supplemented with 0.5 mg/mL Proteinase K) was added to tail or heart tissue samples which were then incubated at 55 °C overnight in a heat block. The following day, samples were mixed thoroughly to ensure complete tissue digestion. Chilled 100% isopropanol (0.5 mL, V/V) was added to each sample, and the mixture was shaken vigorously for 1–2 minutes. Samples were then centrifuged at 13,200 rpm for 10 minutes at 4 °C to pellet the DNA. The supernatant was decanted, and pellet was washed with 0.5 mL of chilled 70% ethanol.

Samples were centrifuged at 13,200 rpm for 5 minutes at 4 °C, and the supernatant was decanted and pellets were dried at 55 °C for 15 minutes. Once dry, 500 µL of resuspension buffer (1 mM Tris 0.01 µM EDTA) was added to each sample followed by vortexing. Samples were incubated in a heat block at 65 °C for at least 1 hour to fully resuspend the DNA. PCR was performed using Qiagen Taq DNA Polymerase (Qiagen) according to manufacturer’s instructions. 32 amplification cycles were used to amplify wild-type (WT), floxed, and recombined DNA. Primers used for amplification are shown in **Table S1.**

### RNA extraction and quantitative PCR

Frozen tissue samples were homogenized in 1 mL Trizol (Invitrogen) reagent using an ultrasonic homogenizer. Samples were centrifuged at 10,000 relative centrifugal force (RCF) for 10 minutes at 4 °C to remove tissue debris. Chloroform (200 µL) was added to each sample, vortexed for 15 seconds, and allowed to incubate on ice for 10 minutes. Samples were centrifuged at 10,000 RCF for 15 minutes at 4 °C. After centrifugation, the upper aqueous phase was carefully transferred to a new 1.5 mL microcentrifuge tube. RNA was precipitated using cold isopropanol following by a 70% ethanol wash step. RNA was resuspended in Diethyl pyrocarbonate (DEPC) water and cDNA libraries were generated using the High-Capacity cDNA Reverse Transcription kit (Applied Biosystems) according to the manufacturers protocol. Quantitative RT-PCR (qPCR) reactions were run using the Sybr Green Master Mix (Applied Biosystems) according to the manufacturers protocol. Cycling was performed on a QuantStudio Real-Time PCR System using SYBR Green detection. Primers used for qPCR are listed in **Table S1**.

### Western blot

Whole heart ventricle tissue or cultured cells were lysed in RIPA buffer (200 mM NaCl, 20 mM Tris, 2 mM EDTA, 1% triton-X, 0.1% SDS) supplemented with protease inhibitor cocktail (1:100, Sigma) and phosphatase inhibitors 2 and 3 (1:100, Sigma). Lysates were centrifuged at 10,000 RCF for 10 minutes to clear lysates and the supernatant was transferred to a new tube. BCA Protein Assay (Pierce) was performed on lysates according to manufacturer’s instructions to determine protein concentration and equal amounts of protein (4 µg) were resolved on a 4-12% SDS-PAGE gel at 120V for 1.5 hours. Following electrophoresis, proteins were transferred onto PVDF membranes (Immobilon-P, Millipore) using a cold-wet transfer system at 200 mA for 2 hours. Membranes were blocked with LI-COR blocking buffer (LI-COR) supplemented with 5% goat serum and 0.1% Tween-20 (TBST) for 1 hour at 22 °C. Membranes were then incubated overnight at 4 °C with primary antibodies diluted in 5% goat serum and LI-COR buffer. Antibodies and concentrations used for western blot can be found in **Table S1.** After washing three times with TBST, membranes were incubated with an infrared secondary antibody (LI-COR). Band intensities were quantified using LI-COR software or ImageJ. GAPDH was used as the loading control for protein abundance normalization.

### Tissue fixation and processing

For histological experiments, mouse hearts were extracted and washed in PBS to remove remaining blood. Hearts were then transferred to 500 mM potassium chloride to arrest hearts in diastole. Hearts were fixed in 4% paraformaldehyde (PFA) for 24 hours, then transferred to 70% ethanol for at least 48 hours, and subsequently embedded in paraffin blocks. Hearts were embedded in a short axis orientation to obtain a 2-chamber view. Tissue sections 4 µm thick were collected every 500 µm starting at the apex.

### Immunostaining and microscopy

PFA-fixed and paraffin-embedded tissue sections underwent deparaffinization with xylene followed by rehydration using a series of ethanol concentrations (100%, 95%, and 75%), followed by a water rehydration. Antigen retrieval was carried out in a pH 6.0 citrate buffer (Sigma-Aldrich C9999) in a water bath maintained at sub-boiling temperature for 30 minutes. Tissue sections were then blocked with 5% normal donkey serum (Sigma Aldrich D9663), 5% BSA (Sigma Aldrich A3294), and 0.1% Triton X (Sigma Aldrich X100) in PBS (PBS-T). Primary antibody incubation was conducted overnight at 4 °C, followed by secondary antibody incubation for one hour at 22 °C. Background fluorescence was mitigated using 0.03% Sudan black in 70% ethanol. For Ethynyl-2’-deoxyuridine (EdU) staining, Alexa flour (AF) 488 ClickIT kit was used according to manufacturer’s instructions. Tissue sections were then incubated in 4’,6-diamidino-2-phenylindole (DAPI) at a concentration of 1:10,000 in PBS. Detailed information regarding primary and secondary antibodies used and concentrations can be found in **Table S1**.

For analysis of cell cycle markers in tissue sections, seven images of the left ventricle were captured per animal at 1000 µm above the apex in P21 mice. These metrics were also assessed after experimental MI in adult mice at 28 dpi. Seven images adjacent to the scar (border zone, BZ) and four images in the remote to the scar (remote zone, RZ) at 20X magnification were taken 1500 µm above the apex per mouse. Imaging was performed on a Nikon NiE Eclipse microscope. All ICD stains were imaged using an inverted Nikon A1 Eclipse Ti Laser Scanning Confocal Microscope equipped with a 60X/1.40 and 100X/1.45 oil immersion objective. Seven images of the left ventricle were captured at 1000 µm above apex per animal.

WGA staining was used for CM cross sectional area measurement. Quantifications were performed on Image J using an algorithm that normalized pixels to µm^2^, rolling ball background, converted positive fluorescent signal and negative background into a binary format, and quantified number of particles per field and cross-sectional area of each particle. Analyze Particles was used with parameters: size (100–∞ pixel²), circularity (0.2–1.0), exclusion of edge artifacts, and inclusion of holes. The output generates a numbered list of measured cells with area values, allowing manual exclusion of misoriented cells or artifacts.

### Immunofluorescence staining of isolated CMs

P21 mouse hearts were digested with 1 mg/mL collagenase type II (Worthington Biochemical) via Langendorff retroaortic perfusion, as previously described.^24,25^ Isolated CMs were washed with PBS and resuspended in 2× microtubule stabilizing buffer (100 mM 1,4-piperazinediethanesulfonic acid, pH 6.9, 30% glycerol, 1 mM EGTA, and 1 mM MgSO4),^26^ followed by the addition of an equal volume of 8% paraformaldehyde prepared in microtubule stabilizing buffer, resulting in fixation with 4% paraformaldehyde for 15 minutes at room temperature. Fixed cells were centrifuged at 300 RCF for 2 minutes, the supernatant was discarded, and the pellet was resuspended in 1 mL PBS.

Cells were then permeabilized and blocked with 0.1% Triton X-100 in 3% bovine serum albumin (BSA, Sigma-Aldrich) at room temperature for 1 hour. Primary antibodies were applied overnight at 4°C in 3% BSA buffer. After thorough washing, the cells were incubated with secondary antibodies (1:400 in PBS) and and/or phalloidin (Invitrogen, A12379) for one hour at room temperature. Cell nuclei were stained with 1 μg/mL DAPI (Sigma-Aldrich) following additional PBS washes. Finally, cells were mounted using ProLong Diamond Antifade Mountant (Thermo Fisher Scientific). Images were obtained using a Nikon A1 laser scanning confocal microscope and the perinuclear microtubule mean fluorescence intensity (within a 1 μm perinuclear region) was analyzed with Fiji software. The distance between the thin filaments (100 filaments per sample, n = 3 per group) was quantified by an add-on module, PatternJ, in ImageJ software.^27,28^

### Electron microscopy (EM)

Tissues were processed for EM by standard protocol. Tissue pieces of approximately 1 mm³ were excised and briefly rinsed with 1mL cold buffer (2.5% glutaraldehyde solution in 0.1 M sodium cacodylate buffer, pH 7.3). Subsequently, the tissues were submerged in 2.5% glutaraldehyde fixative for 1 hour at 22 °C using 2 mL of fixative per vial. After aspiration of the fixative, the tissue blocks underwent three 10-minute washes with buffer. Post-fixation was then carried out with 1% osmium tetroxide in distilled water on ice for 1 hour. Following post-fixation, the tissues were rinsed three times with distilled water for 5 minutes. Dehydration was performed in a series of steps: 70% methanol for 15 minutes, 90% methanol for 15 minutes, and 100% methanol three times for 20 minutes each, followed by two rounds of 15-minute incubations in acetonitrile (or propylene oxide). Subsequently, the tissues were immersed in a 50/50 mixture of epoxy resin and acetonitrile (or propylene oxide) for 1 hour on a rotator, followed by additional incubations in fresh epoxy resin for 1 hour and 2 hours on a rotator, respectively. The tissue samples were then blocked out into labeled molds and polymerized overnight at 70 °C for a minimum of 10 hours. 70 nm Ultrathin sections were cut on an RMC ultramicrotome, mounted on 200 mesh copper grids, stained with a saturated solution of uranyl acetate in 25% methanol and counterstained with Reynolds lead citrate by standard methods. Stained sections were examined in a JEOL 1400 Flash transmission electron microscope operating at 80 KvA. Images were recorded using a Nanosprint12 digital camera operating on AMT image acquisition and analysis software.

### Ventricular CM single cell suspension and nucleation/ploidy analysis

Analysis of CM nucleation and ploidy was performed as described previously.^25^ Juvenile or adult mouse hearts were digested with 1 mg/ml collagenase type II (Worthington Biochemical) via Langendorff retroaortic perfusion. Following perfusion, the atria was excised, leaving only ventricular tissue, which was triturated in Kruftbrühe (KB) buffer, filtered through a 250 µm mesh followed by fixation in 2% PFA at 22 °C for 15 minutes. Cells were blocked in 10% normal donkey serum (Invitrogen) and 0.1% triton-X for one hour at 22 °C on a rocker. Fixed cell suspensions were stained with alpha actinin (1:300) overnight at 4 °C. Alexa flour (AF) 488 ClickIT kit was used to stain for EdU incorporation, AF555 Goat anti-mouse (1:500 was used to detect alpha-actinin. AF647 Goat anti-rabbit (1:500) was used to detect N-cadherin to ensure we are counting intact CMs and DAPI. Cells were washed and spun down. 20 µl of prolong gold (Invitrogen P36930) was added per 10 µl of cells. 30 µl of cells mixed with prolong gold was then pipetted across a superfrost charged slide in a horizontal line and imaging was performed on a Nikon Ti80 microscope.

Slides were systematically scanned using a standard fluorescent microscope to identify EdU-positive (EdU^+^) CMs. During manual scanning, the nucleation status of each labeled cell (i.e., mono-, bi-, tri-, or tetra-nucleated) was recorded, and representative images were captured at 10X magnification. Our goal was to image approximately 60-80 EdU^+^ cells for analysis. To assess DNA content, the fluorescence intensity of any DNA dye within a single nucleus was quantified from the captured images. Fluorescence intensity serves as a proxy for DNA content and was normalized to a reference nucleus of known diploid, such as those from tri- and tetranucleated CMs. Ploidy analysis was performed using NIS Elements BR (version 5.30.05, 64-bit) By integrating nuclear number quantifications from direct microscopic observations with ploidy assessments from image analysis, an estimate of total DNA content per cell was obtained. When ploidy analysis was combined with EdU incorporation, it allowed for the inference of whether cell division had occurred. This method has been described in detail ^25^.

### Adult MI Model

Experimental MI was surgically induced by permanent ligation of the left anterior descending (LAD) artery in adult mice (8-10 weeks) as described previously ^29^. Briefly, mice were anesthetized using a 2% isoflurane and 98% oxygen gas mixture at a flow rate of 2 liters per minute. Chest hair was removed using Nair hair removal (Church & Dwight), followed by cleaning and sanitization of the skin using iodopovidone and ethanol. A left hemithorax incision was made using surgical scissors, and the 4th intercostal space was bluntly dissected to access the thoracic cavity. Manual pressure was applied to the right hemithorax to exteriorize the heart. The LAD was directly visualized and ligated using prolene 6-0 suture. Subsequently, the heart was allowed to retract into the thorax, while pneumothorax was minimized by applying manual pressure to the sternum, and the skin was sutured with prolene 4-0. Post-surgery, mice received subcutaneous injections of buprenorphine (0.5 mg/mL) at a dose of 4 mL/kg of body weight and 300 μL of 0.9% saline solution and allowed to recover under warm light on a heating pad. On the following day after MI surgery, mice were administered meloxicam oral solution (1.5 mg/mL) at a dose of 7.5 mg/kg.

### Electrocardiography

Mice were anesthetized with a 1.5% isoflurane and 98.5% oxygen mixture at a flow rate of 1 liter per minute for electrocardiography assessments. Three-lead electrocardiograms (ECGs) were recorded in anesthetized mice using a standard small-animal ECG acquisition system (PowerLab/8sp) Mice were placed in a supine position on a temperature-controlled platform maintained at ∼37 °C to prevent hypothermia. Subcutaneous needle electrodes were inserted into the right forelimb (RA), left forelimb (LA), and left hindlimb (LL) to obtain standard limb leads I, II, and III. ECG signals were acquired at a sampling rate of 1 kHz and digitally recorded using Labchart 7 pro (version 7.3.8). To minimize electrical noise, the recording environment was shielded, and electrodes were placed with consistent positioning. Signals were continuously monitored for 2 minutes to ensure stability, and at least 60 seconds to 2 minutes of high-quality, motion artifact-free data were selected for analysis. ECG parameters, including heart rate (HR), ΔRR, P duration, PR interval, QRS duration, and QT interval, were analyzed using Labchart 7 pro (version 7.3.8).

### Echocardiography

Mice were anesthetized with a 1.5% isoflurane and 98.5% oxygen mixture at a flow rate of 1 liter per minute for echocardiography assessments. Echocardiography was conducted at 21 days of age in uninjured control and Llgl^CKO^ mice. For adult MI studies, echocardiography was performed at baseline (day −1 prior to experimental MI), as well as on days 3 and 28 post-MI. Echocardiography was carried out utilizing a VEVO 3100 echography machine (FUJIFILM; VisualSonics) equipped with VEVO3100 imaging software (V 3.2.8.15513) and the MX550D probe. The cardiac apex was aligned with the aorta to obtain longitudinal B-mode images of the left ventricle (LV). LV circumference was delineated, and areas quantified during systole and diastole were utilized to calculate percent ejection fraction (%EF) and cardiac volumes. Subsequently, the MX550D probe was rotated 90° to obtain perpendicular cross-sectional images of the left ventricle at the mid-papillary level. M-mode images were used to measure left ventricular (LV) internal diameters during systole and diastole, which were then used to calculate percent fractional shortening (%FS). All images were analyzed using VEVO LAB software (FUJIFILM; VisualSonics).

### Scar size quantification

Tissue sections were prepared as described above and stained with Gömöri’s trichrome stain. Subsequently, tissue sections were scanned at a resolution of 4000 dots per inch using the super COOLSCAN 9000 (Nikon). Quantification of scar area was conducted in ImageJ employing color thresholding tool to delineate the scar (blue) and viable myocardium (pink/purple). Scar size was quantified and averages from two serial tissue sections per heart; one section collected at the mid-papillary level, and one collected 500 µm below the mid papillary level towards the apex of the heart. Scar size was reported as a percentage of the midline length of the left ventricular circumference, and as percentage of total left ventricular area using the semi-automated MIQuant method as previously described ^30^.

### RNAsequencing

MI or sham surgeries were performed on Myh6^Cre^ or Llgl1^CKO^ 8- to 10-week-old mice. At 4 dpi, left ventricles were collected, homogenized, and RNA collected via the RNAeasy Mini Kit (Qiagen). Equal numbers of males and females were used for each of the four groups with n = 3 collected per genotype and condition. Libraries were prepared using SMART-seq mRNA LP (Takara). A Fragment Analyzer 5200 with an HS NGS kit (Agilent) was used to confirm quality of the RNA and constructed libraries. Library sequencing was performed on a NovaSeq X with a 10B flowcell (Illumina) to achieve a ∼100M paired-end reads. Fastq files were processed using Nextflow v.25.04.4 with the nf-core/rnaseq pipeline version 3.22.0. Key steps in the pipeline include quality control (FastQC v0.12.1), read trimming (Trimgalore v0.6.10), alignment (STAR v2.7.11b), and quantification (salmon v.1.10.3). Alignment was done to the GRCm39 mouse transcriptome. Following quantification, factors of unwanted variation were estimated by RUVSeq (v1.40.0) using the RUVr approach. Quantification files were loaded into DESeq2 (v1.44.0) using tximport (v1.32.0) for differential expression analysis, with one RUVr factor regressed out. Differentially expressed genes were defined by FDR < 10% (Benjamini-Hochberg). Analyses and visualization were completed in R v4.4.2. Data are available from the Gene Expression Omnibus (GEO) under accession number GSE313605.

### Neonatal rat ventricular CM (NRVCM) culture and siRNA knockdown

CMs were isolated from postnatal day 1 (P1) neonatal Sprague Dawley rats (Charles River). Rats were sacrificed by decapitation, hearts were excised, atria were discarded, and ventricles were minced. Minced ventricular tissue was digested using the Miltentyi neonatal heart dissociation kit according to the manufacturer’s instructions (Miltenyi Biotec). Digested tissue was passed through a 70 µM cell strainer and cells were then pelleted by centrifugation at 1000 RCF for 5 minutes at 4 °C, washed with HBSS, and resuspended in Dulbecco’s modified eagle medium 4.5 g l −1 glucose (DMEM, Life Technologies) supplemented with 7.5% FBS and passed through an isotonic percoll gradient to separate CMs from other cell types. Enriched CMs were washed 1x with HBSS, resuspended in DMEM supplemented with 15% fetal bovine serum and plated for downstream experiments. For siRNA transfection experiments, NRVCMs were plated in DMEM medium supplemented with 15% FBS. 24 hours after plating, cell media was changed to 7.5% FBS and CMs were transfected with siRNA according to MISSION siRNA protocol (Sigma-Aldrich) with a final molarity of 100 nM per siRNA. Two separate siRNAs were used against Llgl1 (Sima Aldrich: SASI_Rn02_00268906 and SASI_Rn01_00081640 [Ref seq NM_152844]). Cells were incubated for 48 hours prior to collection or fixation for immunostaining. To confirm knockdown, cells were collected in RIPA buffer 48h after transfection and protein was extracted for western blotting.

### Lentiviral construction and in vitro experiments

Lentivirus constructs were developed to expressed wildtype mouse Llgl1 (NM_001159405.1) or mouse phosphomutant Llgl1 whereby serine residues at position 982, 986, 988, 989, 992 were mutated to alanine residues. Three million 293T cells were plated in 10 cm plates the day before lentivirus packaging. Transfection was prepared by mixing 500 µL per dish of Opti-MEM (from Gibco) with PEImax (3 µgPEI:1µg DNA, from polysciences) in tube and incubated for 5 minutes. 5 µg of lentiviral plasmid (pLV[Exp]-mCherry:T2A:Puro-cTnT>EGFP/3xGS/HA, pLV[Exp]-mCherry:T2A:Puro-cTnT>HA/3xGS/mLlgl1[NM_001159405.2]*(S982A, S986A, S988A, S989A, S992A, pLV[Exp]-mCherry:T2A:Puro-cTnT>HA/3xGS/rLlgl1[NM_001304613.1) along with 3 µg of packaging vector (pCMV-dR8.91 or psPAX2) and 1 µg of envelope vector (pMD2.0G) was mixed with in 500 µL Opti-MEM. Plasmid mix was gradually added to the PEImax solution and incubated for 15 minutes. Fresh media was added to 293T cells and 1 mL of the PEI-DNA mixture was added to each plate of cells. Cells were cultured for viral production for 48 hours. Supernatants were then collected and filtered through a 0.45 µm filter. Lentiviral particles were titrated on cultured NRVCMs to determine optimal multiplicity of infection (MOI).

To determine the optimal multiplicity of infection (MOI) for viral transduction, cells were seeded in 24-well plates at a density of 30,000–40,000 cells per well (6–8 wells per condition) and incubated overnight at 37 °C with 5% CO₂. The following day, the virus was added at varying volumes (3 µL, 10 µL, 30 µL, 100 µL, and 300 µL) in the presence of 5 µg/mL polybrene to enhance transduction efficiency.

Cells were incubated overnight with the virus, after which the medium was replaced with fresh culture medium. After 48 hours, transduction efficiency was assessed under a fluorescence microscope by evaluating the expression of the red fluorescent reporter. The optimal viral volume was determined as the condition in which 100% of cells exhibited fluorescence without visible cytotoxicity. For subsequent experiments, the viral volume corresponding to complete transduction (e.g., 600µL for 250,000 cells on 12 well plate) was used as a reference to maintain consistent MOI across experiments.

### Surface biotinylation and enrichment

Neonatal rat ventricular CMs (NRVCMs) were washed with ice-cold biotinylation buffer [10 mM HEPES, 130 mM NaCl, 2 mM MgSO₄, 1 mM CaCl₂, 5.5 mM glucose, pH 7.8–8.0] at 4 °C and kept on ice for the remainder of the biotinylation procedure to halt protein trafficking. Membrane proteins were biotinylated using the cell-impermeant reagent EZ-Link Sulfo-NHS-SS-Biotin (Thermo Fisher Scientific) at 1 mg/mL for 10 minutes. To eliminate unreacted Sulfo-NHS-SS-Biotin, cells were quenched with 100 mM glycine three times.

Cells were lysed and scraped in RIPA buffer supplemented with a 1× protease inhibitor cocktail and 1 mM PMSF by pipetting. Approximately 40 μL of lysis buffer was used per pellet from a single well of a six-well plate. Following a brief incubation on ice (∼2 minutes), lysates were cleared by centrifugation at 15,000g for 10 minutes at 4 °C, and the supernatants were collected and kept on ice. Protein concentration was determined using a BCA assay.

For the isolation of biotinylated proteins, 30-μL aliquots of streptavidin magnetic beads were washed twice with 1 mL of RIPA buffer before incubation with 360 μg (∼90 μL) of whole-cell lysate for 1 hour at room temperature on a rotator. To facilitate binding, 500 μL of additional RIPA buffer was added to each sample. After incubation, the beads were pelleted using a magnetic rack, and the supernatant (flow-through) was collected and stored on ice for subsequent analysis. The beads were subjected to sequential washes to remove nonspecific binders using 1 mL of each buffer per wash: twice with RIPA buffer, once with 1 M KCl, once with 0.1 M Na₂CO₃, once with 2 M urea in 10 mM Tris-HCl (pH 8.0), and twice more with RIPA buffer.

Biotinylated proteins were eluted from the streptavidin beads by boiling in 30 μL of 3× protein loading buffer supplemented with 2 mM biotin and 20 mM DTT for 10 minutes. The samples were vortexed briefly, cooled on ice, and centrifuged to collect condensation. Beads were separated using a magnetic rack, and the eluates were collected. Samples were subsequently resolved on a 4–20% SDS-PAGE gel for downstream analysis.

### Exclusion criteria

We make every attempt to avoid excluding animals or datapoints from any experimental analysis. Animals were excluded from MI studies if they received inadequate MI injury (>45 %EF at 3 dpi) or excessively large injuries (<10% EF at 3 dpi). Datapoints were excluded if they fall >3 standard deviations (SD) outside of the mean. These exclusion criteria are set before the study.

### Data and Statistical Analysis

All data were collected and analyzed in a blinded and randomized manner. Statistical analysis was performed with GraphPad Prism 8.3 software. All data were assessed for normality and equal variance. Statistical comparisons between two groups were analyzed using unpaired, two-tailed unpaired Student’s *t* test or Mann-Whitney u test when variances were significantly different. Between three or more unrelated groups, the data were analyzed by one-way ANOVA followed by Tukey’s multiple-comparison test. Statistical comparison between groups with two variables were analyzed by two-way ANOVA followed by Tukey’s multiple comparison test. Statistical comparison between 2 groups assessed at different time points was performed by two-way repeated measures ANOVA followed by Šidák’s multiple comparison test or repeated measures mixed-effects ANOVA with Fisher’s LSD test when not all time points were accounted for. Survival curves were analyzed by Mantel-Cox test. Ploidy analysis represented as a stacked bar chart totaling 100% was analyzed by MANOVA, Wilks’ Lambda (**Table S2**). All data are presented as mean ± SE unless specified otherwise.

## RESULTS

### Llgl1 deletion impairs proper establishment of CM ICDs

Llgl1 is known to regulate the maintenance of apical junction complexes and is recruited to the basolateral domain after cell-cell contact-initiated polarization.^6^ While *llgl1* global deletion in zebrafish has been shown to disrupt ICD integrity,^21^ whether or not Llgl1 influences mammalian CM junctional complexes or function is unknown. We first explored expression patterns of Llgl1 in mammalian cardiac tissue over the postnatal time course. Llgl1 is highly expressed in the developing postnatal mouse heart and its expression declines postnatally both at the protein (**Fig. 1A,B**) and RNA level (**Fig. 1C**, RNA expression from published data^31^). This expression pattern correlates with total protein level of ICD proteins including α- and β-catenin, Connexin 43, and N-cadherin which also decline at both the protein (**Fig. 1A, S1A,B**) and RNA (**Fig S1C**) level postnatally. Indeed, Llgl1 protein expression significantly correlates with protein expression of α- and β-catenin and N-cadherin measured between P1 and P21 (**Fig. S1B**). Connexin 43 expression, however, did not show a strong correlation with Llgl1 expression (**Fig S1A,C**). Llgl1 expression corresponds to CM terminal differentiation characterized by maturation of ICD whereby N-cadherin migrates to myocyte termini to mediate the formation of specialized junctional structures,^32^ and is coincident with transition from hyperplastic to hypertrophic CM growth.^33^ While Llgl1 expression is enriched in the early postnatal heart, its expression is maintained into adulthood suggesting a functional role for Llgl1 past early postnatal developmental stages.

**Figure 1.**
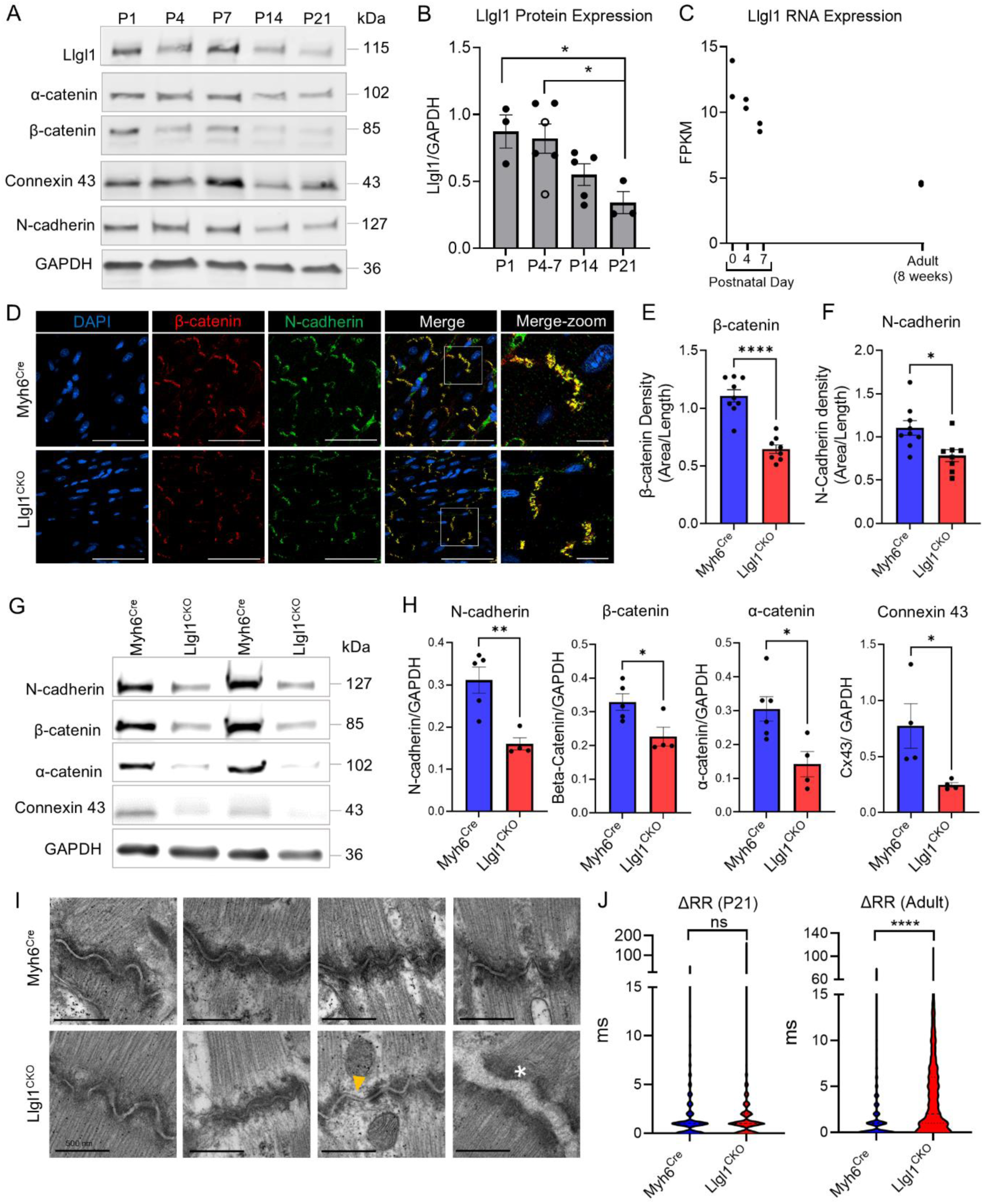
Llgl1 is required for proper ICD establishment in mice. **A)** Expression of Llgl1 and ICD proteins from whole heart ventricle lysates from C57B6/J mice during mouse postnatal development. P = postnatal day. **B)** Quantification of Llgl1 protein expression normalized to GAPDH. n = 3 for P1, 4 for P4 (closed circles), 2 for P7 (open circles), 5 for P14, and 3 for P21. P4 and P7 data were pooled for statistical analysis. One-way ANOVA with Tukey’s test. **C)** RNA expression of Llgl1 from mouse whole heart ventricle samples adapted from ^31^. **D)** Immunohistochemistry of β-catenin and N-cadherin localization in P21 mouse hearts. White box in “Merge” images is shown enlarged in “Merge-Zoom” panels. n = 5 per group. Scale bar = 50 µm and 10 µm for “Merge-zoom” panels. **E)** Quantification of ICD signal, measuring density as the total pixel area divided by length for β-catenin and **F)** N-cadherin. n = 9 Myh6^Cre^ and 7 Llgl1^CKO^ hearts. Unpaired Student’s t-test. **G)** Western blot and **H)** quantification of ICD protein expression from P21 whole heart lysates. n = 5 Myh6^Cre^ and 4 Llgl1^CKO^. Unpaired Student’s t-test and Mann-Whitney test for Connexin 43 assessment. **I)** Electron microscopy from P21 hearts demonstrating decreased protein density (yellow arrow) at the ICD of Llgl1^CKO^ knockout hearts compared to control. White asterisk indicates a CM that appears completely disconnected from an adjacent CM. Yellow arrow denotes a lack of protein density at the ICD. Scale bar = 500 nm. **J)** Delta RR interval collected from electrocardiography measurements of either P21 or adult mice. n = 8 Myh6^Cre^ and 7 Llgl1^CKO^ for P21 electrocardiography and n = 8 Myh6^Cre^ and 7 Llgl1^CKO^ for adult electrocardiography. Unpaired Student’s t-test. * = p < 0.05, ** = p < 0.01, and **** = p < 0.0001, and ns = not significant.

To investigate the role of Llgl1 specifically in mammalian CMs, we crossed Llgl1 floxed (Llgl1^fl/fl^) mice^23^ with Myh6^Cre^ mice to generate a transgenic line whereby Llgl1 is deleted specifically in CMs (Llgl1^fl/fl^;Myh6^Cre^ [Llgl1^CKO^]). PCR of genomic DNA from heart tissue verified proper recombination of the floxed out exon 2 in Llgl1^CKO^ mice shown in lane 1 of **Fig. S1D** (expected size of recombination ∼150 bp), whereas Llgl1^fl/wt^ cre negative animals (**Fig. S1D** lane 2) show the expected non-recombined PCR product of ∼1000 bp and wildtype mice (**Fig. S1D** lane 4) show no PCR band as expected. Quantitative RT-PCR and Western blotting of left ventricular heart tissue from postnatal day 21 (P21) mice confirmed significant reduction of the Llgl1 transcript (**Fig. S1E**) and protein levels (**Fig. S1F,G**) in Llgl1^CKO^ mice compared to littermate controls. Residual Llgl1 expression observed by Western blot is likely due to expression from other cardiac cells, as Llgl1 is not exclusively expressed by CMs in the heart.^34^

Considering the role of Llgl1 in establishing junctional complexes in other cell types, we investigated if Llgl1 is necessary for the development of CM ICDs. In P21 mice, a timepoint when ICDs are fully mature, we found a reduction in the density of several proteins typically localized to the ICD including β-catenin and N-cadherin (**Fig. 1D**) as well as Connexin 43 and α-catenin (**Fig. S2A,C**). We quantified the relative density of these proteins localized to the ICD by measuring the area of protein staining within individual ICDs divided by the length of the ICD and confirmed significantly less protein density for all proteins analyzed (**Fig. 1E,F, Fig. S2B,D**). Consistent with histological findings, total protein expression of N-cadherin, β- and α-catenin, and Connexin 43 was significantly decreased in P21 Llgl1^CKO^ mouse hearts compared to controls (**Fig. 1G,H)**. Electron microscopy revealed overall less electron dense ICDs, and in some cases almost no protein density localized to the ICD in Llgl1^CKO^ mouse hearts as indicated by yellow arrow (**Fig. 1I**). In rare instances we found CMs that appeared completely disconnected from adjacent CMs in Llgl1^CKO^ hearts (white asterisk). ICD morphology was substantially disrupted and a-tubulin staining revealed some apparent microtubule disorganization near the ICDs (**Fig. S3A**). However, we found no overt differences in overall microtubule organization, desmin patterning, or thin filament density at P21 (**Fig. S3A-D**). Collectively, these data demonstrate a critical role for Llgl1 in the proper establishment of CM cell-cell junctions.

ICDs play an instrumental role in a variety of processes including tethering cytoskeletal proteins to the membrane^35^, conduction^36^, and signaling^37^. We investigated the functional consequence of Llgl1 deletion in basal cardiac phenotypes in P21 mice. We found no significant differences in cardiac function (percent fractional shortening [%FS] or percent ejection fraction [%EF]) nor did we find differences in morphological parameters including left ventricular internal diameters [LVID] or volumes during systole [-s] or diastole [-d] or wall thicknesses (LVAW) by echocardiography (**Fig. S4A**). Electrocardiography of P21 Llg1^CKO^ mice revealed no difference in average heart rate (R-R interval) or rate of ventricular repolarization (QTc or JT interval) (**Fig. S4B**), and electrical conduction through the atrioventricular node was also not different between genotypes (P-R interval). However, we observed significantly longer duration of the P-wave and QRS interval in Llg1^CKO^ mice compared to control indicating delayed conduction through the atrial and ventricular tissue, respectively (**Fig. S4B**). Delayed conduction through the cardiac tissue in Llgl1^CKO^ mice is consistent with our observations that Connexin 43 expression and localization is disrupted, as Connexin 43 deletion in mouse CMs has been shown to be associated with QRS complex prolongation.^38,39^ While ventricular conduction was delayed, we found no evidence of ventricular arrhythmias in P21 mice as indicated by deviation in R-R intervals (**Fig. 1J**). However, electrocardiograph analysis of adult mice (12 weeks old) revealed significant deviations in R-R intervals in Llgl1^CKO^ mice compared to Myh6^Cre^ controls (**Fig. 1J, Fig. S5A**), indicating that modulations to ICD integrity contribute to progressive abnormal ventricular rhythm that strongly manifests in adulthood. Llgl1^CKO^ breeders exhibit significantly decreased survival probability compared to Llgl1^fl/wt^;Myh6^Cre^ or Llgl1^wt/wt^;Myh6^Cre^ breeders (**Fig. S5B**), indicating an increased propensity for premature sudden death. Considering observed ventricular arrhythmia in Llgl1^CKO^ mice, we postulate that premature death may be attributed to cardiac conduction defects, however further studies are required to confirm cause of death. Collectively, Llgl1 deletion in CMs impairs proper establishment of ICDs which coincides with cardiac conduction abnormalities and ultimately ventricular arrhythmia.

### Llgl1^CKO^ mice display increased CM cell cycle activity

Llgl1 is considered a tumor suppressor gene in epithelial cells as its downregulation is associated with increased cell proliferation and in some cases tumorigenesis.^40^ CMs are structurally rigid cells and upon maturation become highly resistant to cell cycle activity.^41,42^ Considering CM cell junctions are disrupted in Llgl1^CKO^ mice, we hypothesized that CM cell cycle activity would coincidentally be upregulated in Llgl1^CKO^ mice. We injected Llgl1^CKO^ and littermate control mice with the thymidine analogue EdU at P4 and P5 to track cells undergoing DNA synthesis and collected hearts for histological analysis at P7, a developmental timepoint whereby CM proliferation has mostly ceased in wildtype mice and subsequent heart growth is achieved primarily by hypertrophy of existing CMs.^33^ We performed immunostaining for EdU positive CM nuclei using an anti-Cre antibody to detect CM nuclei (as previously reported^43^) as Myh6^Cre^ and Llgl1^CKO^ mice express Cre specifically in CM nuclei our genetic model. Compared to control, Llgl1^CKO^ animals displayed a significant increase in CM EdU incorporation (**Fig. 2A,B**) as well as increased CM nuclear expression of the cell cycle markers phosphorylated histone H3 (pH3) (**Fig. 2C,D**) and proliferating cell nuclear antigen (PCNA) (**Fig. S6A,B**) at P7. In drosophila, deletion of the Llgl1 homolog, Lgl, results in overgrowth phenotypes thought to be mediated at least in part by increased nuclear localization of the pro-proliferative transcriptional element Yki.^44^ Indeed, at P21 we observed significantly more nuclear localization of the Yki mammalian homolog, Yap, in Llgl1^CKO^ hearts compared to control (**Fig. 2E,F, Fig. S6E**) consistent with observations of increase CM cell cycle activity.

**Figure 2:**
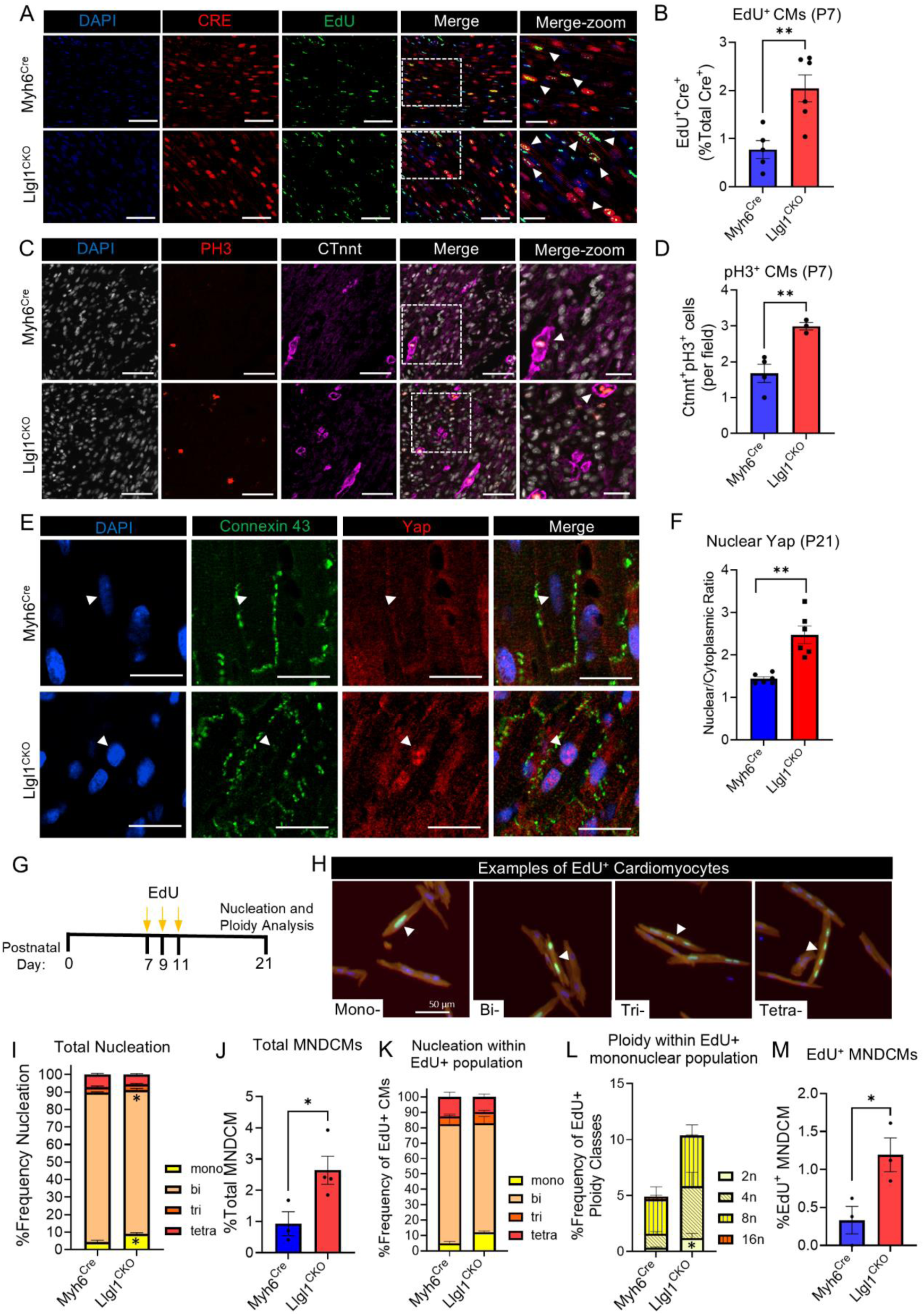
Llgl1^CKO^ juvenile mice display increased evidence of CM cell cycle activity. **A)** Immunohistochemistry staining and **B)** quantification of EdU^+^ CMs from P7 mouse hearts of animals injected with EdU at P4 and P5. n = 6 Myh6^Cre^ and 5 Llgl1^CKO^. Unpaired Student’s t-test. **C)** Immunohistochemistry staining and **D)** quantification of pH3^+^ CMs in P7 mouse hearts. n = 4 Myh6^Cre^ and N=3 Llgl1^CKO^. Unpaired Student’s t-test. **E)** Representative images of Yap nuclear localization in P21 mouse CMs. Arrow denotes CM nuclei. **F)** Quantification of Yap nuclear versus cytoplasmic staining intensity. n = 6 Myh6^Cre^ and 6 Llgl1^CKO^. Mann-Whitney test. **G)** Experimental outline of CM nucleation and ploidy analysis. Yellow arrows indicate single intraperitoneal EdU injection. **H)** Representative examples of EdU^+^ CMs with varying nucleation. White arrow points to EdU^+^ CM. **I)** Distribution of CM nucleation classes (mono- bi- tri- tetra-) including all CMs, both EdU^+^ and EdU^-^. Data are represented at percent out of total CMs counted. **J)** Total MNDCMs considering EdU^+^ and EdU^-^ cells represented as percent of total CMs counted. Unpaired Student’s t-test. **K)** Distribution of CM nucleation classes (mono- bi- tri-tetra-) within the EdU^+^ subset of CMs. **L)** Distribution of ploidy (2N, 4N, 8N, 16N) within the EdU^+^ mononuclear population. MANOVA, Wilks’ Lambda. **M)** Percent of EdU^+^ MNDCMs. Unpaired Student’s t-test. Data for K-M are represented as percent of total EdU^+^ CMs counted. n = 3 Myh6^Cre^ and 3 Llgl1^CKO^ for panels I-M. Dotted box in “Merge” images is shown enlarged in “Merge-Zoom” panels for A and C. Scale bar = 50 µm in first 4 panels for each genotype, scale bar = 20 µm in “Merge-zoom” panels for A and C. Scale bar = 20 µm in L. * = p < 0.05 and ** = p < 0.01.

Expression of cell cycle markers in mature CMs can often be observed in the absence of cell cycle completion, as increased cell cycle activity can result in endomitosis whereby CMs increase nucleation and/or ploidy without cytokinesis.^45,46^ To determine if Llgl1 deletion results in CM cell cycle completion, we administered mice with EdU at P7, P9, and P11, and performed comprehensive CM nucleation and ploidy analysis at P21 (**Fig. 2G,H**). By this analysis, one scenario by which an EdU^+^ CM (a cell that has undergone DNA synthesis) that is both mononuclear and diploid (<u>M</u>ono<u>n</u>uclear and <u>d</u>iploid <u>c</u>ardio<u>m</u>yocytes [MNDCMs)], 2N) is that it arose from a cell cycle completion event.^25^ When considering all CMs (EdU^+^ and EdU^-^) we found more mononuclear cells in Llgl1^CKO^ hearts compared to control (**Fig. 2I**), and upon ploidy analysis of the entire mononuclear population, we found a significant increase in the number of MNDCMs in Llgl1^CKO^ hearts (**Fig. 2J**). The MNDCM population is thought to be more competent to proliferate compared to higher ploidy class CMs^47,48^ supporting the hypothesis that Llgl1 deletion increases CM proliferative capacity. We next evaluated nucleation and ploidy classes within EdU^+^ CMs. Llgl1^CKO^ mice had more mononuclear EdU^+^ CMs overall (**Fig. 2K**), and upon ploidy analysis of the mononuclear population (**Fig. 2L**) we found approximately 3-fold more EdU^+^ MNDCMs in Llgl1^CKO^ hearts (**Fig. 2M**). While observed EdU^+^ MNDCMs could arise from DNA damage,^49^ our collective immunostaining for cell cycle markers and Yap nuclear localization along with analysis of EdU^+^ MNDCMs collectively indicate that Llgl1 deletion increases CM cell cycle activity between P7 and P21.

### Llgl1^CKO^ mice show improved functional recovery following myocardial infarction (MI)

Alterations to CM ICDs are known to mediate CM mechanotransduction, which is activated post MI and mediates pathophysiological phenotypes.^50^ Yap nuclear activity in CMs, for example, improves ejection fraction post MI likely through pro-proliferative or perhaps anti-apoptotic pathways.^51,52^ Considering parallels between phenotypes observe in Llgl1^CKO^ mice and protective or pro-reparative mechanisms in the literature, we next examined whether Llgl1^CKO^ adult mice display improved functional response after cardiac injury. Experimental MI by permanent ligation of the LAD coronary artery or sham operation was induced in 8-week-old mice and animals subjected to MI were administered with daily intraperitoneal injection of EdU from 4 to 10 dpi to track cells undergoing DNA synthesis (**Fig. 3A**).^25^ Both Llgl1^CKO^ and control mice received comparable MI injuries as evidenced by similar %EF at 3 dpi (**Fig. 3B**). While %EF and %FS declined in control animals over the course of the 28-day study, these functional parameters were improved in Llgl1^CKO^ animals compared to Myh6^Cre^ controls (**Fig. 3B** and **Fig. S7A**). Strikingly, Llgl1^CKO^ mice showed substantial resistance to ventricular dilation compared to control which was apparent in both systolic and diastolic measurements at 28 dpi (**Fig. 3C,D Fig. S7B,C,** representative 28 dpi shown in **Fig. 3E**). Sham operated animals showed no significant difference in cardiac function between genotypes over the duration of the 28-day study (**Fig. 3B-D, Fig. S7A-C**). Histological analysis of trichome stained injured hearts revealed significantly smaller fibrotic scar size in Llgl1^CKO^ mice as assessed by both scar length and total scar area (**Fig. 3F-H)**. The left ventricular anterior wall, where majority of CM death occurs following MI, was thicker at the mid-papillary level in Llgl1^CKO^ mice at 28 dpi (**Fig. 3I**) further indicating smaller injury size at 28 dpi in Llgl1^CKO^ mice. CM cross sectional area in both Llgl1^CKO^ sham operated and Llgl1^CKO^ 28 dpi hearts was slightly but significantly larger than compared to control (**Fig. 3 K,L**) implicating morphological CM differences between genotypes under both conditions. Collective echocardiography and histological data indicate that Llgl1^CKO^ mice are protected from adverse ventricular function and remodeling following injury.

**Figure 3:**
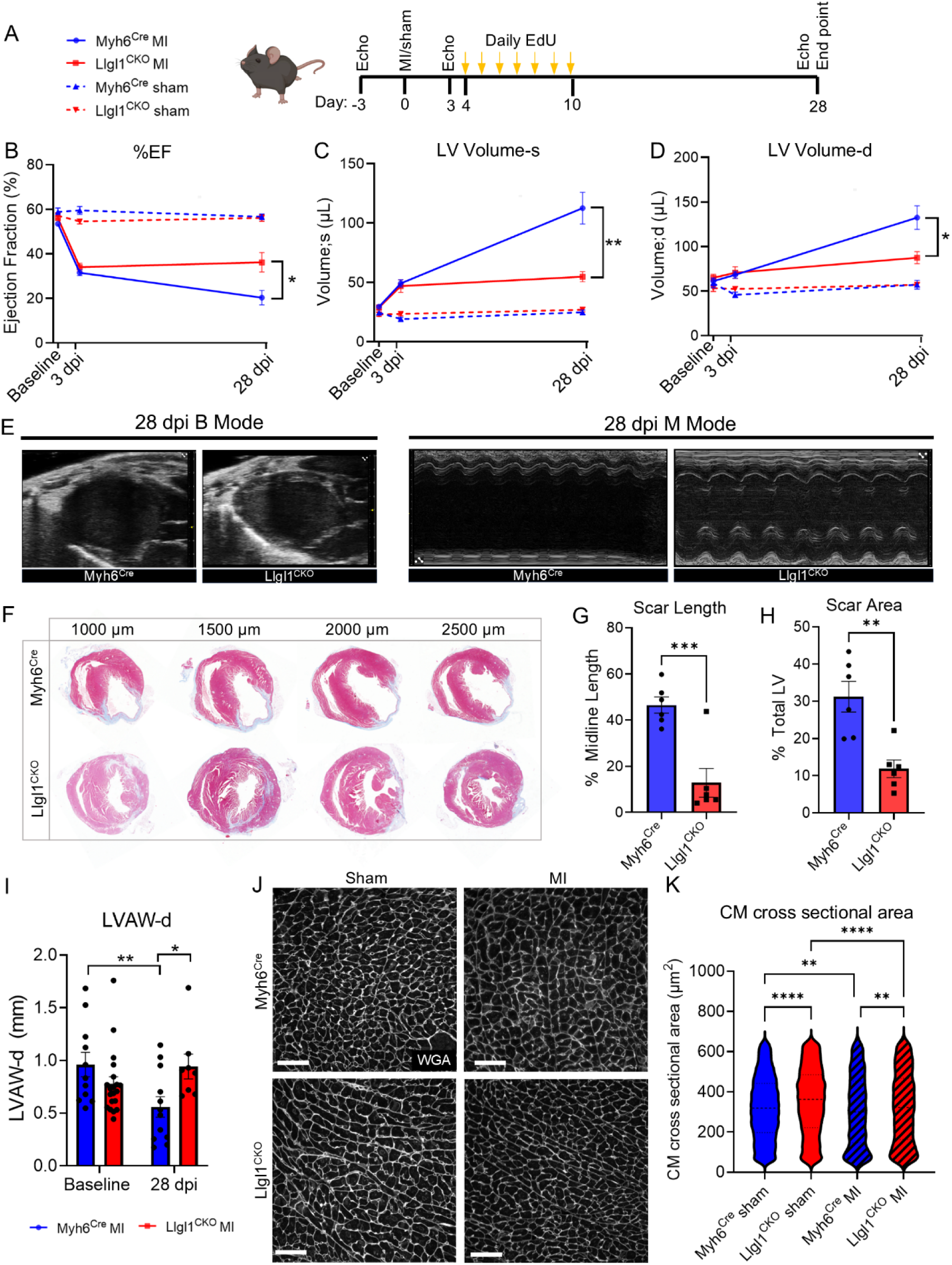
Llgl1^CKO^ mice display improved functional and histological outcomes following experimental MI. **A)** Experimental outline of sham or MI experiments performed on Llgl1^CKO^ and littermate control mice at 8-weeks of age. **B)** Quantification of %EF and **C)** left ventricular (LV) volume during systole and **D)** diastole at baseline (prior to MI) and at 3 and 21 dpi. Solid line indicates MI animals, dotted lines indicate sham operated animals. n = 12 Myh6^Cre^ MI, 8 Llgl1^CKO^ MI, 8 Myh6^Cre^ sham, and 6 Llgl1^CKO^ sham. No significant differences identified between genotypes in sham group. Repeated measures two-way ANOVA with Šidák test. **E)** Representative images of B mode and M mode echocardiography data collected at 28 dpi. **F)** Representative images of Gömöri trichome stained heart sections collected at 28 dpi. **G)** Quantification of scar area calculated as percent midline length and **H)** percent total left ventricular area. n = 5 Myh6^Cre^ and 6 Llgl1^CKO^ for panels G and H. Unpaired Student’s t-test. **I)** Left ventricular anterior wall thickness measured during diastole (LVAW-d) from short axis M mode. n = 12 Myh6^Cre^ baseline, 20 Llgl1^CKO^ baseline, 11 Myh6^Cre^ 28 dpi, and 5 Llgl1^CKO^ 28 dpi. Repeated measures mixed-effects ANOVA with Fisher’s LSD test. **J)** Representative WGA staining and **K)** quantification of CM cross sectional area from hearts of MI and sham operated mice collected at 28 dpi. n = 7 Myh6^Cre^ MI, 6 Llgl1^CKO^ MI, 8 Myh6^Cre^ sham, and 6 Llgl1^CKO^ sham. Two-way ANOVA with Tukey’s test. * p<0.05, **p<0.01, ****p<0.0001.

### Transcriptional analysis of sham operated and post infarct hearts

We next performed RNA-sequencing on sham operated and post infarct hearts at 4 days post operation to further understand potential mechanisms by which Llgl1^CKO^ animals may display improved outcomes following MI. Transcriptional profiles form distinct clusters by genotype in both sham and post MI conditions (**Fig. 4A,B**). We input 356 differentially expressed (DE) genes between Myh6^Cre^ and Llgl1^CKO^ sham operated hearts (**Table S3**) and 362 DE genes between Myh6^Cre^ and Llgl1^CKO^ post infarct hearts (**Table S4**) with false discovery rate of <0.1 into Ingenuity Pathway Analysis. Pathway analysis of DE genes from sham operated hearts revealed a striking upregulation of cell cycle and mitotic genes In Llgl1^CKO^ compared to Myh6^Cre^ control hearts (**Fig. 4C**). Cell cycle related genes were specifically involved in kinetochore assembly and chromatin segregation (*Sgo1*, *Bub1*, *Kntc1*, *Cenpn*)^53,54^ and related to Rho GTPase-Formin signaling, which coordinates actin remodeling and promotes cytokinesis across cell types including CMs (**Fig. 4C,D**).^55,56^ These data support our observation of increased cell cycle activity in uninjured Llgl1^CKO^ mice and suggest that Llgl1 deletion may enable kinetochore assembly and chromosome segregation to facilitate CM cytokinesis.

**Figure 4.**
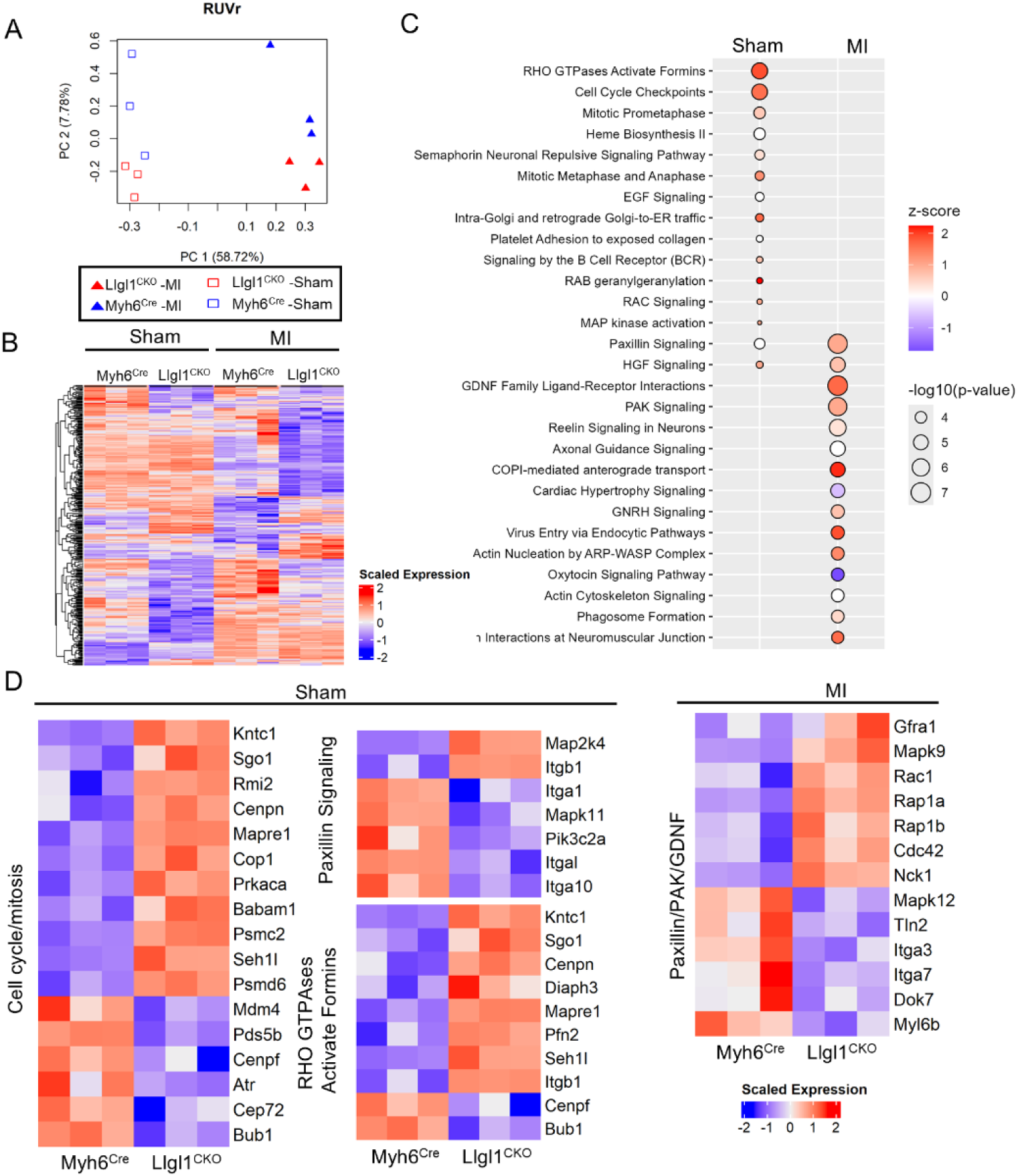
Transcriptional analysis of sham and post MI hearts at 4 days post operation. **A)** PCA plot of transcriptional profiles from 12 RNAseq samples after RUVSeq. **B)** Combined heatmap of all DE genes with FDR <0.1 between Myh6^Cre^ and Llgl1^CKO^ in either sham or MI hearts. **C)** Top enriched pathways of differentially expressed genes between Llg1^CKO^ and Myh6^Cre^ sham (left column) or MI (right column) hearts. Genes with Padj. of <0.1 were used for pathway analysis. **D)** Heatmap of genes comprising identified pathways shown in C.

Paxillin signaling, a key pathway mediating mechanotroansduction was modulated in both Llgl1^CKO^ sham and post infarct hearts compared to Myh6^MCM^ controls (**Fig 4. C,D**). Genes comprising the Paxillin pathway included several integrins (Itga1, *Itgal*, *Itga10*, *Itga3*, *Itga7*) that were downregulated in either sham or post MI Llgl1^CKO^ hearts and *Itgb1* which was strongly upregulated in Llgl1^CKO^ sham hearts. Integrins play a crucial role focal adhesion assembly, and they interact directly with Paxillin to facilitate recruitment of the focal adhesion complex.^57^ Interestingly, Itgb1 has been shown to protect CMs from ischemic injury,^58^ suggesting Itgb1 overexpression may be a potential mechanism by which Llgl1^CKO^ hearts display improved outcomes following MI. In addition to Paxillin signaling, at 4 days post MI Llgl1^CKO^ hearts showed strong enrichment in differentially expressed genes related P21 activated protein kinase (PAK) and GDNF signaling. CM Pak1 signaling is protective following both pressure overload^59^ and cardiac ischemia-reperfusion injury,^60^ again aligning gene expression profiles in Llgl1^CKO^ hearts with pathways that confer protection from cardiac injury. Collectively, these data suggest a state of altered mechanotransduction and cell cycle in Llgl1^CKO^ hearts that may confer protection from ischemic injury. While our data do not definitively rule out contribution from other cell types in the divergent response to MI in Llgl1^CKO^ versus Myh6^Cre^ hearts, we observed minimal differential expression of immune response or fibrotic remodeling genes, for example, at this early post infarct timepoint.

### Improved cardiac function following MI is associated with increased CM cell cycle activity in Llgl1^CKO^ mice

Compromised ICDs persist into adulthood in uninjured Llgl1^CKO^ mice (**Fig. S7E,F**), and RNAsequencing indicates strong upregulation of cell cycle and mitosis gene networks in Llgl1^CKO^ hearts. At 28 dpi following MI, we observed a noticeable decrease in fascia adherens proteins at the ICD in the border zone (**Fig. 5A**). Not only was the protein density of N-cadherin and β-catenin decreased in adult Llgl1^CKO^ hearts post MI (**Fig. 5B**), but the co-localization of these proteins was disrupted as well (**Fig. 5C**). Collectively, these data suggesting CMs maintain a state of altered mechanotransduction and cellular plasticity into adulthood. We next asked if improved functional outcomes following MI in Llgl1^CKO^ mice were associated with increased CM cell cycle activity following injury. EdU was administered daily from day 4-10 post MI, and quantification of EdU^+^ CMs revealed a significant increase in the percentage of EdU^+^ CMs in both the border zone and remote zone of adult Llgl1^CKO^ mice following MI (**Fig. 5D-F**). Furthermore, Llgl1^CKO^ hearts displayed a ∼2.5 fold increase in pH3^+^ CMs at 28 dpi (**Fig. 5G,H**) suggesting increased CM cell cycle activity persists for weeks following MI.

**Figure 5:**
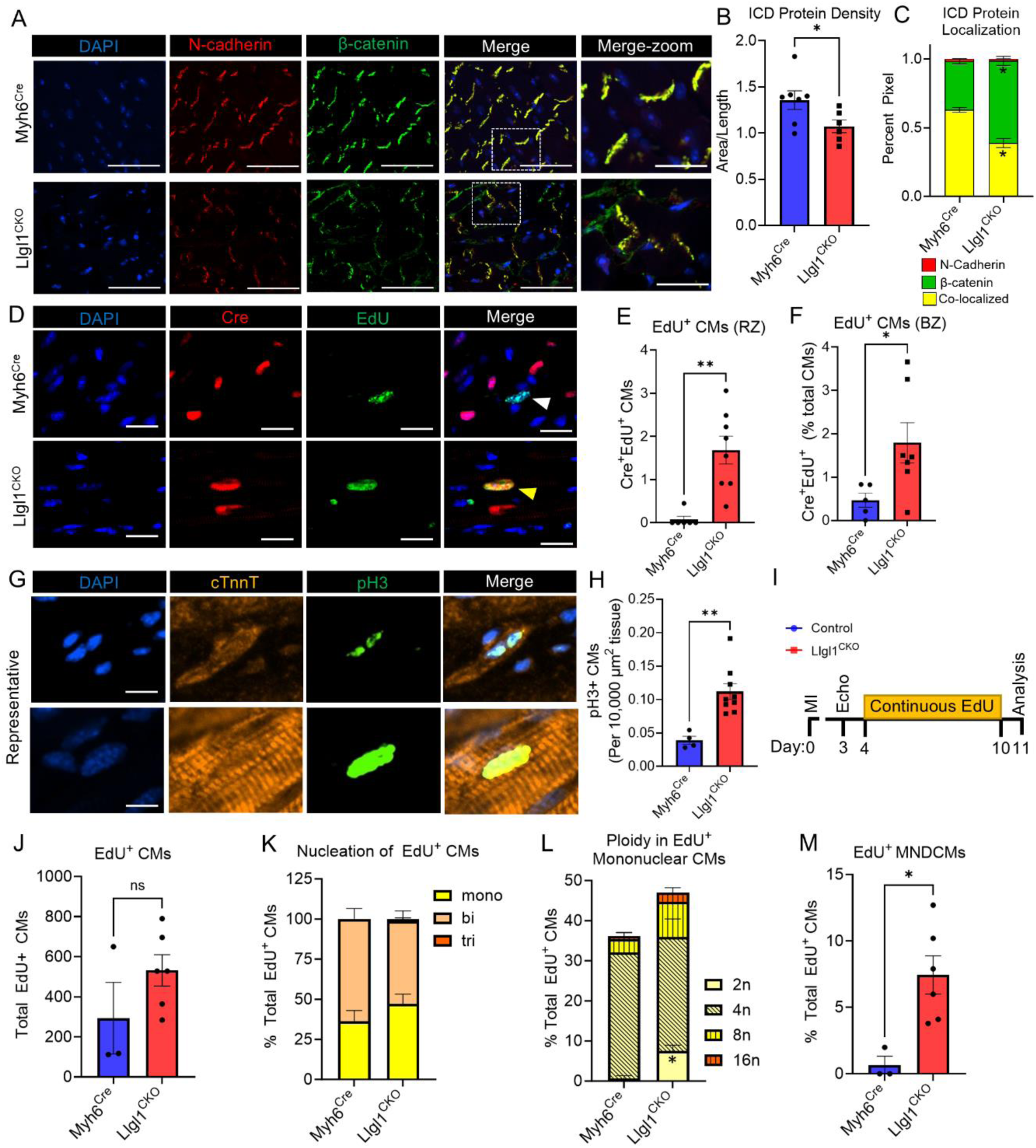
Llgl1^CKO^ mice have increased CM cell cycle completion following MI. **A)** Representative images of N-cadherin and β-catenin immunostaining from 28 dpi infarcted hearts in the remote region. Dotted box in “Merge” images is shown enlarged in “Merge-Zoom” panels. Scale bar = 50 µm and 20 µm in “Merge-zoom” panels. **B)** Quantification of ICD signal, measuring density as the total pixel area divided by length for β-catenin and N-cadherin. n = 7 Myh6^Cre^ and 6 Llgl1^CKO^. Unpaired Student’s t-test. **C)** Quantification of co-localization of N-cadherin and β-catenin (yellow) versus non co-localized β-catenin (green) and N-cadherin (red). n = 7 Myh6^Cre^ and 6 Llgl1^CKO^. Unpaired Student’s t-test. **D)** Representative images of Cre immunostaining and EdU co-staining from 28 dpi infarcted hearts in the border zone region. White arrow indicates EdU^+^ Cre^-^ (non-CM) nucleus. Yellow arrow indicates EdU^+^ Cre^+^ (CM) nucleus. Scale bar = 20 µm. **E)** Quantification of remote zone (RZ) and **F)** border zone (BZ) EdU^+^ CM nuclei as indicated by positive Cre staining. n = 6 Myh6^Cre^ and 8 Llgl1^CKO^. Mann-Whitney test. **G)** Representative images of pH3^+^ cTnnt^+^ cells and **H)** quantification of total pH3+ CMs per 10,000 µM^2^ of cardiac tissue. n = 4 Myh6^Cre^ and 9 Llgl1^CKO^. **F)** Experimental design for assessment CM nucleation and ploidy following MI in 8-week-old mice. **G)** Total number of EdU^+^ CMs found in each preparation. Mann-Whitney test. **H)** Distribution of CM nucleation classes (mono- bi- tri-) within the EdU^+^ subset of CMs. **I)** Distribution of ploidy (2N, 4N, 8N, 16N) within the EdU^+^ mononuclear population. MANOVA, Wilks’ Lambda. **J)** Percent of EdU^+^ MNDCMs. Data for H-J are represented as percent of total EdU^+^ CMs counted. n = 3 Myh6^Cre^ and 6 Llgl1^CKO^ in panels G-J. Mann-Whitney test. * = p < 0.05, ** = p < 0.01, and ns = not significant.

We again rationalized that increased CM EdU incorporation and pH3 expression in Llgl1^CKO^ hearts could either be indicative of CM cell cycle completion or could indicate an endomitotic event in the absence of cytokinesis (i.e. further polyploidization). To test if adult post infarct CMs were completing the cell cycle, we conducted nucleation and ploidy analysis on isolated EdU^+^ CMs from adult mice following MI. We performed LAD ligation on 8-week-old mice and at 4 days post operation we implanted osmotic minipumps for continuous delivery of EdU. After one week, we removed the osmotic minipumps and allowed a 24-hour chase period to allow cells to complete cytokinesis (**Fig. 5I**). We then digested hearts into single cell suspensions and performed comprehensive nucleation and ploidy analysis on CMs. We found a trend towards increased total EdU^+^ CMs in Llgl1^CKO^ hearts (**Fig. 5J**), consistent with our histological findings, although this difference did not reach statistical significance likely due to high variability in this parameter and inefficient digestion of border zone cells. While total EdU^+^ mononuclear CMs were slightly expanded in Llgl1^CKO^ hearts (**Fig. 5K**) we observed a nearly eight fold expansion of the EdU^+^ MNDCM population when analyzed as a percent of total EdU^+^ CMs (**Fig. 5L,M**). Considering there is a trend towards increased total number of EdU^+^ CMs in Llgl1^CKO^ hearts, the actual fold increase in CMs completing cytokinesis in Llgl1^CKO^ hearts is likely substantially greater than eight-fold. Nevertheless, our data indicate that Llgl1 deletion in CMs increases CM cell cycle activity and cytokinesis in post infarct adult hearts, which is associated with improved functional outcomes.

### Inducible Llgl1 deletion in adulthood results in improved outcomes post MI

We next asked if protection from myocardial injury in Llgl1^CKO^ mice was a result of permanent deletion of Llgl1 in CMs throughout development, or if Llgl1 deletion around the time of injury could confer similar protection. We performed MI surgeries on adult conditional knockout mice whereby Llgl1 is deleted in CMs following tamoxifen induction (Llgl1^TamCKO^) or control mice carrying tamoxifen inducible cre under the Myh6 promoter (Myh6^MCM^). Tamoxifen was administered 2-3 days prior to MI surgery (**Fig. 6A**) followed by echocardiography and end point histological analysis. We confirmed proper exon recombination by PCR of genomic DNA from hearts of a separate cohort of Llgl1^fl/fl^;Myh6^MCM^ mice 7 days after treatment with two daily intraperitoneal doses of tamoxifen (**Fig. S8A**). There was no significant difference in post MI mortality between Llgl1^TamCKO^ and Myh6^MCM^ mice (**Fig. 6B**) but interestingly, Llgl1^TamCKO^ mice showed significant improvements in %EF at both 3 and 28 dpi (**Fig. 6C**). We observed trends towards improved %FS and reduced ventricular volumes and diameters (**Fig. 6D,E, Fig. S8B-D**), however these did not reach statistical significance.

**Figure 6:**
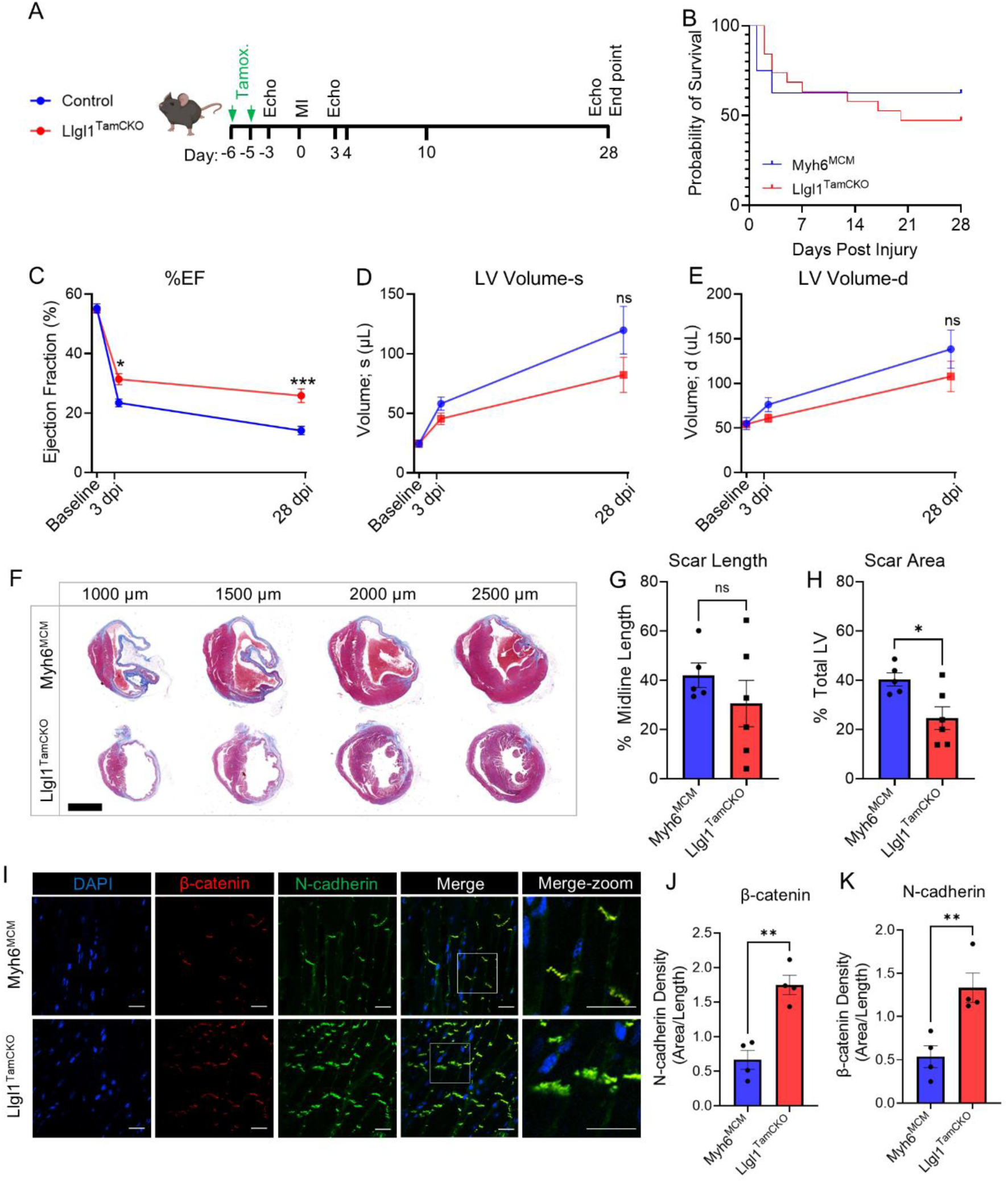
CM deletion of Llgl1 in adulthood protects against adverse remodeling post MI. **A)** Experimental outline of MI experiments performed on Llgl1^TamCKO^ and Myh6^MCM^ mice at 8-weeks of age. **B)** Survival probability of mice post MI. n = 8 Myh6^MCM^ and 19 Llgl1^TamCKO^. Mantel-Cox test. **C)** Quantification of %EF and **D)** left ventricular volume during systole and **E)** diastole at baseline (prior to MI) and at 3 and 21 dpi. n = 5 Myh6^MCM^ and 8 Llgl1^TamCKO^. Repeated measures two-way ANOVA with Šidák test. **F)** Representative images of Gömöri trichome stained heart sections collected at 28 dpi. Scale bar = 2 mm. **G)** Quantification of scar area calculated as percent midline length and **H)** percent total left ventricular area. n = 5 Myh6^MCM^ and 6 Llgl1^TamCKO^. Mann-Data in G was assessed by Whitney test and unpaired Student’s t-test for H. **I)** Immunohistochemistry of β-catenin and N-cadherin localization in adult mouse hearts. White box in “Merge” images is shown enlarged in “Merge-Zoom” panels. n = 5 per group. Scale bar = 20 µm. **J)** Quantification of ICD signal, measuring density as the total pixel area divided by length for β-catenin and **K)** N-cadherin. n = 4 Myh6^MCM^ and 4 Llgl1^TamCKO^ hearts. Unpaired Student’s t-test. * = p < 0.05, ** = p < 0.01, *** = p < 0.001, and ns = not significant.

Llgl1^TamCKO^ hearts showed no significant difference in scar length (**Fig. 6F,G**), but a significant decrease in scar area (**Fig. 6F,H**) further illustrating Llgl1 CM deletion in adulthood protects from adverse outcomes post MI. Llgl1^TamCKO^ ICDs at 28 dpi appeared disrupted compared to Myh6^MCM^ as indicated by β-catenin and N-cadherin immunostaining (**Fig. 6I**). Interestingly and unlike Llgl1^CKO^ hearts, the density of β-catenin and N-cadherin immunostaining was increased in Llgl1^TamCKO^ hearts (**Fig. 6J,K**), perhaps due to endocytic recycling failure of N-cadherin that has already been localized to ICDs in adult Llgl1^TamCKO^ hearts prior to Llgl1 deletion, as Llgl1 has been shown to be involved in N-cadherin endocytic recycling.^6^ Nevertheless, our data demonstrate that both Llgl1^CKO^ and Llgl1^TamCKO^ display disruptions in ICD morphology and protection from MI injury albeit ICD phenotypes appears distinct depending on developmental timepoint whereby Llgl1 is deleted from CMs.

### Llgl1 phosphorylation at the C-terminus domain is required for efficient N-cadherin membrane localization

Our *in vivo* data clearly demonstrate that ICD development is disrupted in Llgl1^CKO^ mice, and impaired ICD development is associated with sustained CM cell cycle activity and improved functional outcomes following injury. Our data from P21 uninjured hearts demonstrate that ICD proteins are not only improperly localized but total abundance of ICD proteins in whole heart lysates is also decreased (**Fig. 1**). Mis-localization of ICD proteins such as N-cadherin, for example, subsequently results in its degradation,^18^ so we developed *in vitro* assays to test if Llgl1 directly influences proper membrane localization of ICD proteins. We found that siRNA mediated Llgl1 knockdown (**Fig. S9A,B**) in cultured NRVCMs resulted in decreased protein density of N-cadherin and Connexin 43 localized to the cell membrane (**Fig. 7A,B, Fib. S9E,F**), and we observed particularly striking enrichment of Connexin 43 localized to the perinuclear space (**Fig. S9E,F**). We repeated these experiments with other ICD proteins such as β-catenin, however, we did not find consistent or convincing staining patterns of β-catenin in NRVCMs suggesting this culture system is not an ideal model for localization of all ICD proteins. Total protein levels of Connexin 43 and N-cadherin were not different between the groups (**Fig. S9A-D**) suggesting that Llgl1 plays a primary role in ICD protein localization as opposed to overall protein expression.

**Figure 7.**
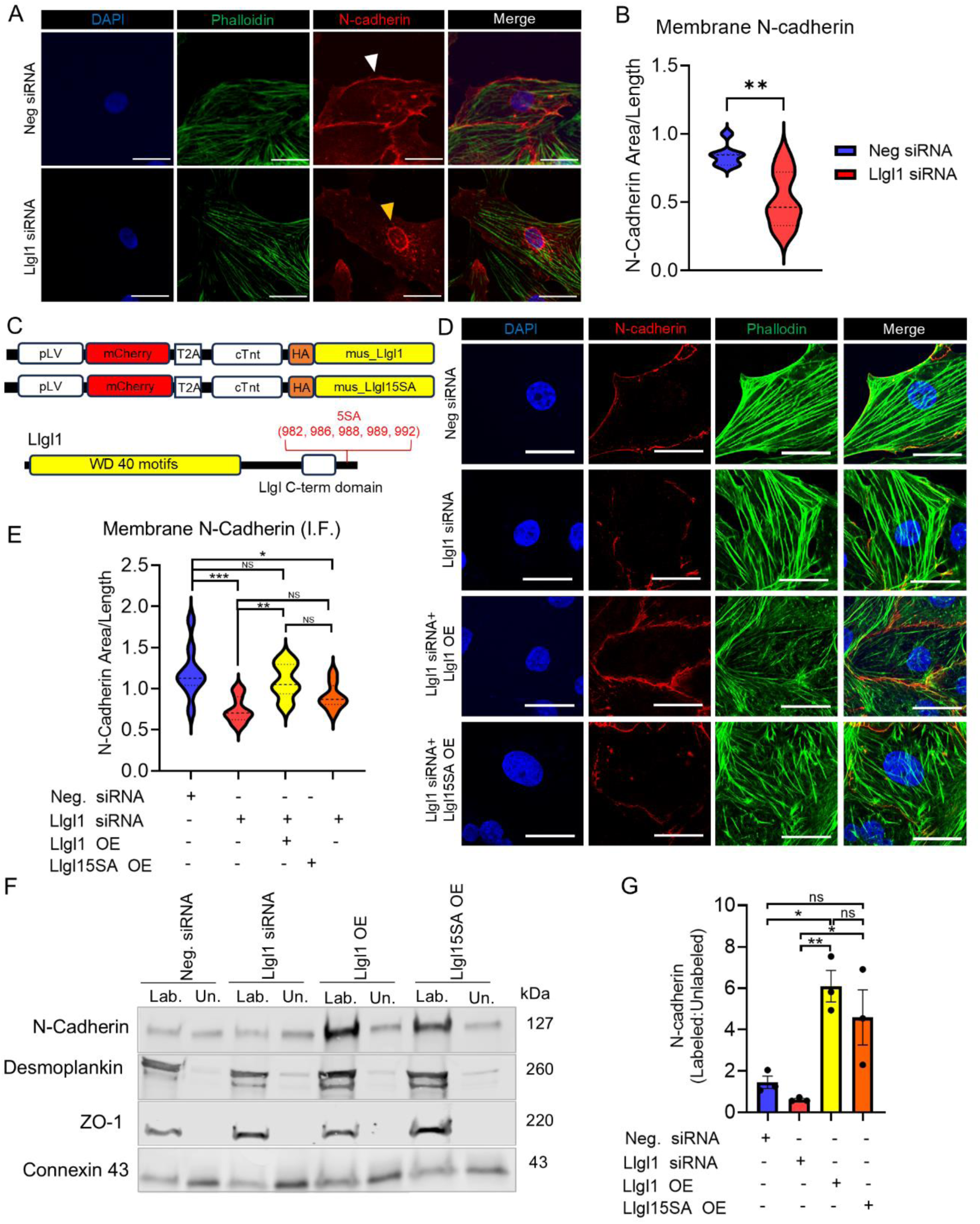
Llgl1 is required for proper membrane localization of N-cadherin in NRVCMs. **A)** Representative immunofluorescent staining of phalloidin and N-cadherin in NRVCMs treated with scrambled or siRNAs against Llgl1. Scale bare = 20 µm **B)** Quantification of N-cadherin membrane localization corresponding to panel A. Values are represented as area of membrane staining divided by the length of entire membrane. Neg = scrambled negative control. n = 7 Neg and 8 Llgl1 siRNA. Unpaired Student’s t-test. **C)** Schematic representation of lentiviral expression constructs (top) and position of S residues mutated to Alanine in the Llgl1 5SA construct (bottom). **D)** Immunostaining of NRVCMs treated with scrambled or Llgl1 siRNAs along with lentivirus over expressing wildtype Llgl1 or Llgl15SA. Scale bar = 20 µm. **E)** Quantification of membrane localized N-cadherin related shown in panel D. Values are represented as area of membrane staining divided by the length of entire membrane. n = 9 Neg, 10 Llgl1 siRNA, 10 Llgl1 OE, and 8 Llgl15SA OE. One-way ANOVA with Tukey’s test. **F)** Western blot of surface biotin immunoprecipitated lysates from NRVCMs treated with scrambled siRNA, Llgl1 siRNAs, or with lentivirus over-expressing wildtype Llgl1 or Llgl15SA. Lab. = biotin labeled fraction, Un. = unlabeled fraction. **G)** Quantification of N-cadherin Western blot protein density related to panel F. Values represented as ratio of labeled to unlabeled fraction for each treatment group. n = 3 per group. One-way ANOVA with Tukey’s test. * = p < 0.05, ** = p < 0.01, *** = p < 0.001, and ns = not significant.

Llgl1 activity is known to be regulated by phosphorylation whereby aPKC directly phosphorylates Llgl1 to alter its affinity with the Par complex.^6,61^ While phosphorylation of serine residues by aPKC at residues S654, S658, and S662^7,62^ has been well established to alter Llgl1 localization, a recent phospho-proteomics experiment of regenerating neonatal mouse hearts found Llgl1 to be differentially phosphorylated at 5 serine residues at its C terminus (S982, S986, S988, S989, and S992),^22^ which are distinct from the classically described aPKC target residues. This cluster of Serine residues, particularly S982 and S986 are highly conserved across mammalian species (**Fig. S9G**). We next tested the in vitro effect of wildtype or phosphomutant Llgl1, whereby Serine residues at positions 982, 986, 988, 989, and 992 are mutated to Alanine (Llgl15SA, [**Fig. 7C**]), by lentiviral transduction (Construct design represented in **Fig. 7C**) in NRVCMs. Lentiviral mediated expression of wildtype Llgl1 fully rescued membrane localization of N-cadherin in NRVCMs following Llgl1 siRNA knockdown. Compared to wildtype, overexpression of the Llgl1 phospho-mutant, displayed attenuated ability to localize N-cadherin to the cell membrane as indicated by N-cadherin immunostaining (**Fig. 7D,E**). Surface biotinylation confirmed that lentiviral mediated overexpression of Llgl1 promoted surface localization of N-cadherin compared to non-transduced NRVCMs, whereas overexpression of Llgl15SA had an attenuated effect on N-cadherin membrane localization (**Fig. 7F,G**). Interestingly, the effect of Llgl1 overexpression appeared to be somewhat specific to N-cadherin, as we observed no difference in surface localization of other membrane proteins including desmoplankin, ZO-1, and Connexin 43. These data indicate that phosphorylation of Llgl1 at the C-terminus of the protein is required for optimizing Llgl1 function in mediating N-cadherin membrane localization in CMs.

## DISCUSSION

Llgl was among the first tumor suppressor genes identified, with early studies in *Drosophila* linking zygotic *lgl* mutations to a hyperproliferative “giant larva” phenotype.^3^ These findings established loss of cell polarity as a hallmark of cancer, underscoring its association with uncontrolled proliferation and disrupted cell-cell junctions. In various cellular contexts, cell-cell junctions transcend their structural roles to dynamically regulate processes critical for tissue homeostasis, such as communication, differentiation, migration, and proliferation. These properties are especially relevant in the heart, where the ICD not only ensures mechanical and electrical coupling but also imposes structural rigidity that is associated with CM terminal differentiation and cell cycle exit.^16^ This rigidity, while essential for synchronized contraction, may constrain cellular plasticity that would otherwise facilitate cardiac regeneration.

Cell polarity is intimately interconnected with development of cell-cell adhesion complexes. In epithelial cells for example, Lgl1 is known to be required for proper formation of apical junctional complexes which are structures that physically distinguish the apical and basolateral membranes.^63,64^ During cardiac morphogenesis, junctional proteins including catenins, connexins, and N-cadherin are localized along the circumferential border of CMs. As CMs elongate during development, these junctional proteins migrate to the CM termini and form the mature ICD.^65^ Our study reveals a critical role for Llgl1 in regulating the proper localization of ICD proteins in CMs. In Llgl1^CKO^ hearts we observed reduced ICD protein density and mis-localization of adhesion molecules which was associated with ventricular arrythmia. These data suggest that Llgl1 plays a conserved role in regulating formation of junctional complexes across cell types including CMs.

While ICD integrity and protein localization is globally disrupted in Llgl1^CKO^ mice, our *in vitro* data suggest that Llgl1 specifically drives membrane localization of N-cadherin in CMs. N-cadherin, a component of the adherens junction in CMs, plays a pivotal role in overall ICD structural maturation.^18,32^ Conditional deletion of N-cadherin in mouse CMs completely disrupts not only adherens junctions but also results in global dissolution of the ICD, indicating that proper N-cadherin localization is a fundamental requirement for ICD development.^18^ The factors that regulate proper localization of N-cadherin are not well understood, but our data indicate that Llgl1 expression is a key factor that regulates membrane localization of N-cadherin in CMs. Based on existing literature and our data we postulate that a primary insult in Llgl1^CKO^ mice is failure to localize N-cadherin to the ICD which subsequently results in global disruption of ICDs.

Our study reveals an association between compromised ICD integrity and increased CM cell cycle activity. The link between cell junctions and proliferation has been well established in the cancer literature whereby loss of polarity and cell adhesions is associated with exacerbated cell proliferation.^4^ Beyond its role in maintaining electrical and mechanical synchronicity, the ICD also has been described as a hub for proliferative signaling in CMs. For example, CM deletion of the junction scaffold proteins ZO-1 and ZO-2 increases CM proliferation by inducing nuclear localization of the pro-proliferative transcriptional regulator Yap.^66^ Similarly, CM deletion of αE- and αT-catenin, proteins that directly bind N-cadherin at the adherens junctions, also results in increased total CM number and greater CM proliferation postnatally. αE and αT-catenin deletion sequesters Yap to the adherens junctions and when deleted, Yap is free to enter the nucleus and drive CM proliferation.^67^ These collective studies illustrate a complex and influential role for ICD proteins in altering CM proliferation throughout development and postnatally. Enhanced nuclear localization of Yap, or perhaps modulation of other mechanotransduction components such as Paxillin or integrin signaling, due to ICD instability may contribute to the enhanced CM cell cycle phenotypes observed in Llgl1^CKO^ hearts.

The dual roles of Llgl1 in regulating both structural and signaling components of the ICD emphasize its significance in maintaining CM integrity. By imposing polarity and stability at cell junctions Llgl1 may limit CM plasticity, a feature critically influential during the post injury response. Indeed, we observe improved cardiac functional outcomes post MI in Llgl1^CKO^ mice which was associated with increased CM cell cycle activity. While the total number of cycling CMs was increased in Llgl1^CKO^ hearts, this number as a percent of total CMs was relatively small. We postulate that improved functional outcomes may be attributed to increased cell plasticity or altered expression of PAK, Paxillin or integrin signaling pathways that have been shown to protect CMs from infarct injury.^58,60^ These collectively pathways along with enhanced CM cell cycle activity could all collectively contribute to improving cardiac function and attenuating post MI ventricular dilation. Our data aligns with the hypothesis that loss of cell polarity—often observed in various proliferative disorders—permits reactivation of cell division by loosening rigid cell junctions, altering cellular geometry and promoting a more permissive environment for cell cycle re-entry. Overall, our data illustrate a key role for Llgl1 in maintaining ICD integrity during development and into adulthood and link disruption of ICDs with CM cell cycle activity and protection from adverse remodeling post injury.

## NOVELTY AND SIGNIFICANCE

### What is known?

- Llgl1 is a polarity protein that is required for proper cell junction formation and that suppresses proliferation in *Drosophila* and mammalian cell types such as epithelial cells.
- Proper localization of N-cadherin to the CM termini is known to be a key requirement of ICD development, yet the comprehensive mechanisms regulated N-cadherin localization remain unknown.

### What new information does this article contribute?

- Llgl1 genetic deletion disrupts proper establishment of ICD proteins and permits CM cell cycle activity into adulthood, and improves functional outcomes following MI.
- Llgl1 is involved in proper membrane localization of N-cadherin in CMs, and phosphorylation of Llgl1 at its C terminus is required for driving efficient N-cadherin localization.

## ACKNOWLEDGEMENTS

We acknowledge the Medical College of Wisconsin Proteomic Core and the Children’s Hospital of Wisconsin Histology Core. We thank Dr. Valeri Vasioukhin at the Fred Hutchinson Cancer Research Center, Seattle Washington for providing Llgl1^fl/fl^ mice and Dr. Sarah J. Parker at Cedar Sinai for consultation on proteomic experiments. This work was supported by the NIH (HL170547 and HL170547-S1 to CCO and BAL, HL141159 and HL141159-S1 to CCO, HL155085 and HL 155085-04S1 to MP, 1R01HL171224 to LH, HL180515-01 to CL, support on T32-HL134643 to ABN), the Medical College of Wisconsin Cardiovascular Center Cullen Healthy Heart Scholar Award (to CCO), the American Heart Association (23SCEFIA1156132 to CL), and the Advancing a Healthier Wisconsin Endowment (to CL). Echocardiography was performed on the High-Frequency Ultrasound Imaging System supported by the NIH S10ODO25038.

## Non-standard Abbreviations and Acronyms

CM: cardiomyocyte
ICD: intercalated disc
CKO: conditional knockout
Tam: Tamoxifen
P: postnatal day
BZ: border zone
RZ: remote zone
dpi: days post injury
MNDCM: mononuclear and diploid cardiomyocyte

